# Expanding the chicken gut virome: Uncovering viral diversity, host interactions, and regional variations across the intestinal tract

**DOI:** 10.1101/2025.04.15.647674

**Authors:** Ji-Xin Zhao, Hong-Bo Ni, Fu-Long Nan, Qiulong Yan, Shenghui Li, Jin-Wen Su, Yanan Cai, Jin-Xin Meng, Hai-Long Yu, Kai-Meng Shang, Rui Liu, Xin-Wen Hou, Hany M. Elsheikha, He Ma, Xiao-Xuan Zhang

## Abstract

Chicken gastrointestinal virome comprises a complex and diverse viral community with significant implications for host health and microbiome function. We analyzed 3,312 publicly available chicken gut metagenomic datasets to establish the chicken gastrointestinal virome collection (CGVD), which includes 39,380 non-redundant viral operational taxonomic units (vOTUs); notably, 84.90% (33,433/39,380) represent novel sequences absent from current databases. Over half of the CGVD vOTUs were classified as bacteriophages, predominantly from the order Caudovirales. The predicted hosts were mainly prokaryotes, particularly Bacillota and Bacteroidota, revealing a multifaceted landscape of virus-host interactions. Many vOTUs infected multiple bacterial phyla, indicating high adaptability and broad ecological impact. In addition, lifestyle prediction showed that 28.28% (11,137 /39,380) of the vOTUs in CGVD were identified as lytic phages. Functional annotation demonstrated that viral genes contribute to key metabolic processes, including nucleotide and amino acid metabolism, thereby facilitating viral replication and host adaptation. The detection of auxiliary metabolic genes and carbohydrate-active enzymes underscores the role of viruses in modulating the gut microbiome. Although antibiotic resistance genes and mobile genetic elements were present, their contribution to horizontal gene transfer appears limited. Additionally, marked regional differences in virome composition were observed between the small and large intestines, particularly in the abundance of families such as Siphoviridae and Myoviridae. CGVD not only highlights the key role of viruses in shaping the chicken gut microbiome and influencing microbial dynamics and metabolic pathways, but also provides new resources and insights for future research.

## Introduction

Chickens are among the most important agricultural species worldwide, serving as a major source of meat and eggs. Their health plays a critical role in ensuring food security and agricultural productivity [1-3]. A key aspect of chicken health lies in the gastrointestinal (GI) tract, which houses a complex and dynamic microbial community. This community is vital for processes such as digestion, nutrient absorption, immune regulation, and overall health [4]. Within this intricate ecosystem, microorganisms—including bacteria, fungi, and viruses—work together to regulate immune responses, combat pathogens, and maintain host health. Therefore, maintaining the homeostasis of the intestinal microbiota is essential for the well-being of chickens [5].

Among these microorganisms, bacteriophages (viruses that specifically infect and lyse bacteria) have emerged as pivotal regulators of microbial populations and functions in the gut [6]. By preying on bacteria and facilitating horizontal gene transfer, bacteriophages significantly influence the composition and genetic diversity of bacterial communities [7]. As such, a comprehensive analysis of the chicken GI virome is essential for promoting chicken health and improving production efficiency.

However, our current understanding of the chicken GI virome remains limited. Existing studies often rely on datasets that are neither large enough nor sufficiently detailed to capture the virome’s full complexity. For example, the chicken multi-kingdom microbiome catalog contains only about 3,000 high-quality viral genomes [8]. While this dataset has advanced our knowledge of the chicken enteric virome, it falls short of encompassing the full diversity of viruses present in the chicken gut. To address this gap, establishing the world’s largest and most comprehensive chicken GI virome database is of immense scientific value. Such an effort would enable a systematic analysis of viral diversity and distribution across different intestinal segments, including the duodenum, jejunum, ileum, cecum, and colon This, in tur, is critical for improving disease prevention, vaccine development, and the formulation of effective antiviral strategies.

Recent advancement in high-throughput sequencing technologies have made it possible to extract viral genomes from metagenomic data, enabling the establishment of large viral genome collections [9]. Examples include the human enteric viral genome collection [10], the ruminant enteric bacteriophage collection [11], and the porcine enteric viral collection [12]. These collections have provided valuable data for studying enteric viruses while offering methodological insights for developing similar resources for chickens. Building on this foundation, constructing a comprehensive chicken GI virome database will significantly enhance our understanding of enteric viral diversity and functions. It will also provide robust support for health management, virus surveillance, and vaccine development in poultry.

In this study, we assembled 3,312 publicly available metagenomic datasets from 18 countries to establish an expanded catalog of the chicken GI virome. This new resource, named the Chicken Gastrointestinal Virus Genome Catalog (CGVD), incudes 39,380 non-redundant viral genomes. These genomes were curated by removing viruses with nucleotide similarity exceeding95%. Remarkably, 33,433 of these viral genomes are novel, not found in any existing databases.

Using the CGVD, we conducted an in-depth analysis of chicken gastrointestinal viruses, exploring their classification, host range, lifestyle, and functional characteristics. Our study also provided the first systematic examination of viral distribution and diversity across different intestinal regions. The CGVD catalog sets a new benchmark in virome research, with implications that extend from basic science to practical applications in poultry health and productivity. The data from this research offer strong support for future initiatives aimed at detecting intestinal diseases of chicken and developing effective antiviral prevention and control strategies.

## Methods

### Chicken metagenomic sample collection

We conducted a comprehensive search of the NCBI database up to September 2023, focusing on Chicken GI tract metagenomic samples annotated with terms such as Chicken*, *Gallus gallus*\*, and related keywords. This search resulted in the identification of 3,312 chicken GI tract metagenomic samples derived from 49 distinct studies (Table S1).

### Preprocessing and assembly

To ensure data quality, the collected samples underwent rigorous preprocessing using Fastp [13] with the parameters -u 30 -q 20 -l 60 -y --trim_poly_g. This step removed adapters and low-quality sequences. The reads that matched the Host genomic sequences were filtered out by aligning reads to the *Gallus gallus* reference genome (RefSeq assembly: GCF_016699485.2) using bowtie2 v2.4.4 [14]. Reads that did not match the host genome were retained as clean reads. These clean reads were subsequently de novo assembled into contigs for each sample using MEGAHIT v1.2.9 [15] with the parameters --k-list 21,41,61,81,101,121. This process generated over 4.1 million long contigs, with a minimum length of 2 kbp and a total length of 38.3 Gbp, with 38.9% of these contigs exceeded 5 kbp in length.

### Identification of viral sequences

We analyzed viral sequences within chicken GI tract metagenomic samples based on established methods [16, 17]. Assembled contigs longer than 5000 bp were selected for virus identification. These contigs were first assessed using CheckV v1.0.1 [18], excluding those with a host gene count exceeding 10 or five times the number of viral genes. Provirus fragments were also identified and using CheckV [18]. The remaining contigs were considered potential viral sequences if they met any of the following criteria: 1) a higher number of viral genes compared to host genes, as determined by CheckV v1.0.1 [18] classification as viral sequences by DeepVirFinder v1.019 with thresholds of score >0.90 and *p*-value <0.01 [19]; and 3) identification as viruses using VIBRANT v1.2.1 [20] with default parameters.

To minimize contamination from non-viral sequences, we applied BUSCOs [21] to detect bacterial universal single-copy orthologs within the potential viral genomes using hmmsearch with default parameters. For each genome, we calculated the BUSCO ratio (number of BUSCOs divided by the total gene count). Genomes with a BUSCO ratio ≥5%, were excluded from further analysis. This process yielded 1,891,878 potential viral genomes. Quality assessment using CheckV [18] classified these genomes as follows: 0.94% (17,870) complete viruses, 1.42% (26,944) high-completeness viruses, 3.55% (67,139) medium-completeness viruses, 50.26% (950,847) low-completeness viruses, and 43.82% (829,078) undetermined completeness viruses. For downstream analysis, we selected viral genomes classified as medium quality or higher based on CheckV quality assessments.

### Viral clustering and gene calling

To de-replicate viral sequences, we employed the following steps: 1) all 111,716 viral sequences (Table S3) were aligned in pairs using BLASTn v2.9.0 [22] with the options -evalue 1e-10-word_size 20-num_alignments 99999. 2) viral sequences sharing ≥95% nucleotide identity across ≥70% of their length were grouped into viral operational taxonomic unit (vOTUs) using custom scripts. 3) For each vOTU, the longest sequence within each vOTU was designated as the representative sequence for further analysis.

This process resulted in 37,020 vOTU sequences derived from all chicken GI tract samples, which were integrated into the Chicken Gastrointestinal Virome Database (CMGC). To evaluate overlap with other virome databases, we applied the same clustering approach to e databases, such as IMG/VR v4 [23], GVD [24], GPD [25], MGV [26], PVD [27], URPC [28], and CMKMC [8]. During this process, 2,360 medium- or higher-quality viruses unique to the CMKMC database were added to the collection. Ultimately, a non-redundant viral genome dataset of 39,380 chicken GI tract viruses with a quality score >50% was constructed (Table S4).

To organize the 39,380 vOTUs into genus- and family-level groups, we performed additional clustering steps: We used the Prodigal v2.6.3 [29] with the -p meta option to predict 2,020,371 putative protein sequences within the vOTUs. Pairwise protein sequence alignments were then performed using DIAMOND v2.0.6.144 [30] with the options -e 1e-5 –max-target-seqs 99999. Subsequently, we calculated the percentage of shared genes and the average amino acid identity (AAI) between vOTU pairs. Using criteria from Nayfach et al [26], we retained vOTU connections with >20% AAI and >10% shared genes for the family level clustering. At the genus level clustering, we retained vOTU connections with >50% AAI and >20% shared genes. Finally, clustering was performed based on vOTU connections using MCL with the -I 1.2 option for family-level clustering and -I 2 for genus-level clustering. This hierarchical clustering grouped the vOTUs in the CGVD into approximately 403 family-level and 3,085 genus-level groups.

### Viral taxonomy

Viral sequences were classified by comparing their protein sequences against a comprehensive reference database. This database included proteins from the Virus-Host DB (downloaded in May 2024), crAss-like proteins from Guerin’s study [31], and viral proteins identified in studies by [32]. Protein-coding sequences of the viral genomes were predicted using Prodigal v2.6.3 [29] with the parameter -p meta. The predicted proteins were then compared to the combined database using diamond v2.0.13 [30], with the parameters --id 30 --subject-cover 50 --query-cover 50 --min-score 50. Classification criteria were adjusted based on the genome size. For small viral genomes (fewer than 30 genes), classification into a known viral family required that more than one-fifth of their proteins matched proteins from that family. For large viral genomes (30 or more genes), at least 10 their proteins needed to match the same viral family [16]. This approach ensured a robust and accurate classification of viral sequences across varying genome sizes.

### Virus host and lifestyle prediction

We performed host and lifestyle predictions of viruses in the CGVD catalog because they can offer valuable insights into the ecological roles, adaptability, and functional contributions of the chicken gut virome. Virus host prediction was conducted using a comprehensive dataset comprising 25,825 metagenome-assembled Genomes (MAGs) collected from various sources. This dataset included 12,339 MAGs from the National Microbiological Data Center (NMDC, https://nmdc.cn/icrggc/), 6,087 MAGs from the Figshare repository (DOI: http://doi.org/10.6084/m9.figshare.24681096.v1). In addition, the 1,450 broiler genomes (1,389 MAGs and 61 isolated genomes) and 5,949 laying chicken MAGs, both collected from our laboratory, are accessible in the Zenodo repository under accession ID 14432035 (https://doi.org/10.5281/zenodo.14432035) and under accession number PRJNA1099794, respectively. Host prediction was performed by analyzing the chicken intestinal prokaryotic genome catalog using two approaches: CRISPR-spacer matches and prophage blasts.

For CRISPR-spacer matches, CRISPR spacer sequences were first predicted using MinCED [33] v0.4.2 with the parameter -minNR 2. Viral genomes were then assigned to a host if the spacer sequence from the host matched the viral genome with a bit-score ≥ 45. These matches were identified using BLASTn with the parameters -evalue 1e-5-word_size 8-num_alignments 999999 [24]. For prophage blasts, viral sequences were aligned against host genome sequences. A host was assigned if the viral sequence matched the host genome at ≥ 90% nucleotide identity and ≥ 30% viral coverage [24].

To classify the lifestyles of vOTUs in the CMGC, we employed DeePhage v1.0 [34]. This tool uses deep neural networks to classify short contigs from metagenomic data by analyzing features of phage DNA and protein sequences. Based on the DeePhage scoring system, vOTUs were categorized into four lifestyle groups: temperate (score ≤ 0.3), uncertain temperate (score 0.3-0.5), uncertain virulent (score 0.5-0.7), and virulent (score >0.7). Higher scores indicated greater, virulence.

### Phylogenetic analysis

Phylogenetic analysis was performed on complete vOTUs, with quality assessment performed using the CheckV algorithm. Following established practices from previous studies, a whole-genome similarity-based phylogenetic approach was employed. To classify previously uncharacterized viral genomes, a viral proteome tree was generated for the complete vOTUs in the CGVD using ViPTreeGen v1.1.2 [35], a tool designed for this purpose. The resulting proteome tree was then visualized using iTOL v6.3.2 [36], providing an intuitive representation of viral relationships and classifications.

### Functional annotation

To elucidate the functional role of predicted proteins, we compared protein-coding genes against several key databases: KEGG [37], Pfam [38], and VOG [39], using VIBRANT [20]. This comparison allowed for the identification of viral auxiliary metabolic genes (AMGs). In addition, we searched the CAZy [40], Virulence Factor Database [41], and a customized antimicrobial resistance database [16] using Diamond v2.0.6.144 [30].

For CAZy annotations, a protein was considered successfully assigned to a specific KEGG functional ortholog or CAZyme if it achieved a score of at least 60 across 50% of the sequence length. For annotations from the Virulence Factor Database, a protein was assigned to a specific virulence factor if it met a threshold of 60% identity across at least 50% of the sequence length.

For antimicrobial resistance gene annotation, we applied specific criteria based on previous studies: (1) beta-lactamase-associated proteins were annotated with a minimum amino acid similarity of 90%, (2) proteins related to multidrug resistance were annotated with a minimum amino acid similarity of 70%, and (3) other antimicrobial resistance genes were annotated using a minimum similarity threshold of 80% [42]. Finally, predicted genes were identified as mobile genetic element (MGE)--like genes by searching open reading frames (ORFs) against a custom MGE database, as previously described [43]. This research was conducted using BLASTn with parameters of an e-value ≤ 10^-5, coverage ≥ 80%, and identity ≥ 70%.

### Spatial profiling of the chicken GI virome across intestinal segments

To investigate the spatial distribution of the chicken GI virome, we analyzed all vOTUs in the CGVC across metagenomes from five distinct intestinal segments: duodenum, jejunum, ileum, cecum, and colorectum This analysis encompassed 575 deeply sequenced metagenomes, each containing over 10 million clean reads (Table S2). Clean reads from each metagenomic sample were aligned to the vOTUs in the CGVC using Bowtie2 [44] with the following parameters: –end-to-end –fast –no-head –no-unal -u 10000000 –no-sq. The abundance of each vOTU was determined by aggregating the number of reads mapped to it. These read counts were subsequently normalized to transcripts per million (TPM), yielding a “read abundance” metric for each vOTU. The relative abundance of a vOTU was calculated by normalizing its abundance relative to the total number of reads mapped in each sample. For family-level analysis, the relative abundances of vOTUs within the same family were aggregated to generate a summary profile.

### Statistical analysis and visualization

All statistical analyses were performed using R version 4.2.2. the Shanno and Richness indices were employed to obtain the relative abundance data of viruses in each sample. Diversity and richness across different groups were assessed using the Wilcoxon rank-sum test and Fisher’s test. To assess beta diversity, principal coordinate analysis (PCoA) was performed using Bray-Curtis distance, and statistical significance was determined with permutational multivariate analysis of variance (PERMANOVA). The Wilcoxon rank-sum test was used to evaluate statistically significant differences in diversity indices and the relative abundance of taxa and functional gene signatures between groups. Sparse curves were generated using the vegan R package (v2.6-4), with chord diagrams were visualized using the circlize R package (v2.8.0). All other visualizations were produced using the ggplot2 package (v4.2.3)

## Results

### Construction of a catalog of chicken GI viruses

To address the challenge of characterizing chicken GI virome, we compiled a comprehensive dataset consisting of 3,312 publicly available metagenomic datasets from the chicken GI tract. This represents the largest chicken GI metagenomic collection to date, spanning 49 studies across multiple regions: 3 Asian countries (China, India and Singapore), 2 North American countries (United States and Canada), 11 European countries (Austria, Belgium, Bulgaria, Denmark, France, Germany, Italy, Netherlands, Poland, Spain and United Kingdom), 1 Oceania country (Australia), and 1 African country (Gambia) (Fig. 1A).

**Figure 1.**
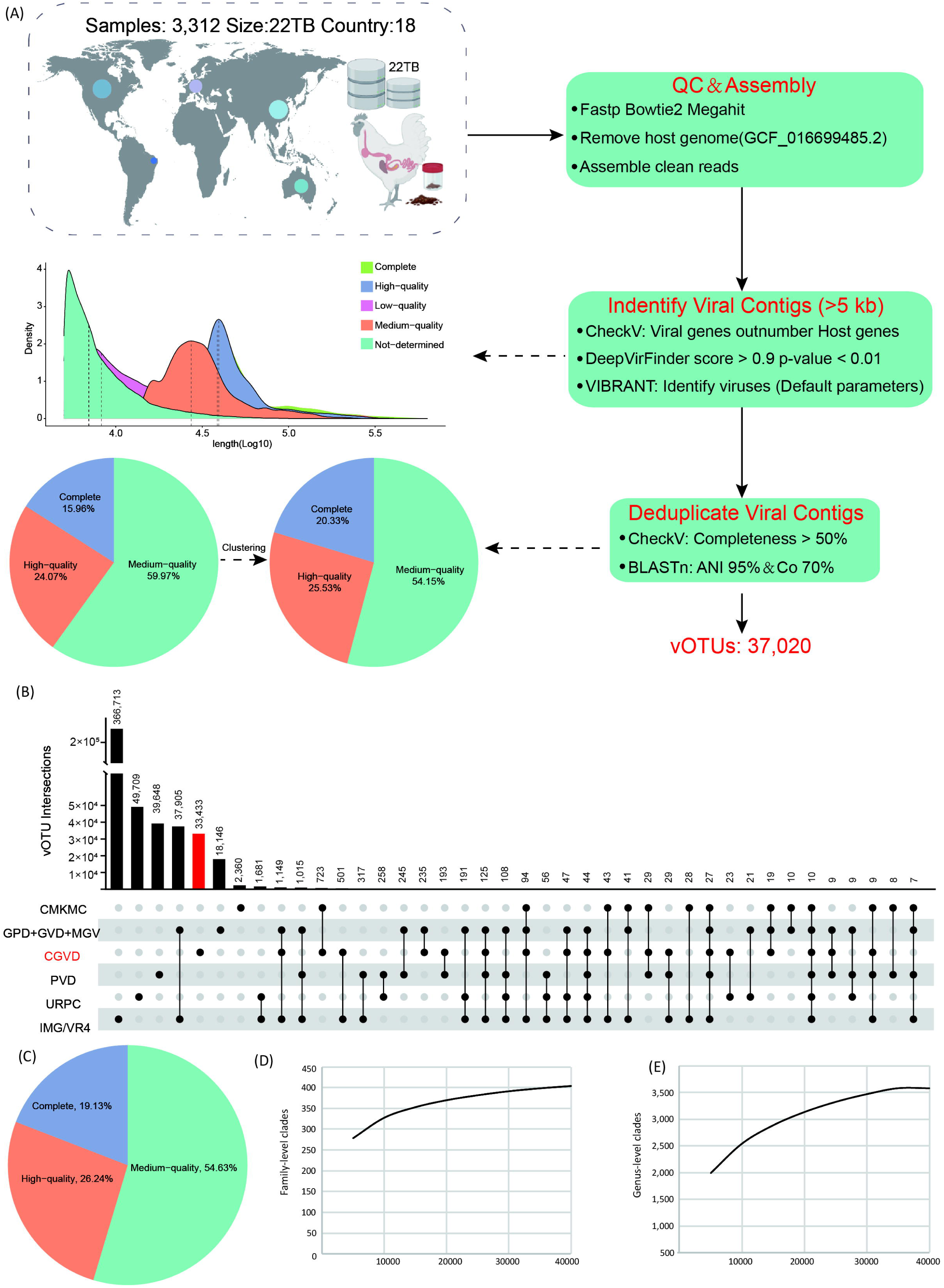
(A) Pipeline for viral Genome extraction and characterization from metagenomic data. (B) UpSet plot illustrating the overlap of vOTUs shared among existing virome databases, including the Chicken Multi-Kingdom Microbiome Catalog (CMKMC), Human Enteric Virome Database (GVD), Human Enteric Phage Database (GPD), Human Metagenomic Enterovirus Database (MGV), Pig Virome Database (PVD), Unified Ruminant Phage Catalogue (URPC), and the Expanded Database of Uncultivated Virus Genomes (IMG/VR4). (C) Quality assessment of viruses included in the CGVD catalog. (D) Cumulative curves representing the growth of approximately family-level groups within the CGVD catalog. (E) Cumulative curves representing the growth of approximately genus-level groups within the CGVD catalog.

To ensure the accuracy and comparability of viral genomes in CGVD with other established datasets [45, 46], we applied stringent quality control measures during data processing. These included the removal of host genomic sequences and comprehensive quality evaluations using CheckV. This approach revealed that the CGVD contains 22 terabytes (Tb) of high-quality reads and over 15.98 million contigs (≥5 kb) (Table S1), demonstrating the robustness of our assembly pipeline. The data processing minimized contamination and low-quality sequences, making the CGVD a reliable resource for future metagenomic studies and ensuring the precision of downstream analyses. Using a combined homology- and feature-based pipeline (the detailed process is described in the methods), we identified approximately 2.1% (111,936) of these contigs as highly credible viral sequences. These viral sequences were then clustered at a threshold of >95% nucleotide similarity [47] to produce a non-redundant CGVD comprising 37,020 vOTUs (Fig. 1A). The vOTUs ranged in length from 5,000 bp to 635,137 bp, with an average length of 38,562 bp and an N50 length of 43,918bp (Table S4).

Low-quality sequences from viral datasets in IMG/VR4, MGV, GVD, GPD, URPC, PVD, and CMKMC were excluded, retaining vOTUs with genome fragments of greater than 50% completeness. These vOTUs were then compared with those in the aforementioned viral genome catalogs. Remarkably, 90.31% (33,462 out of 37,020) of CGVD genomes were unique and did not cluster with any vOTUs from these other datasets. Even against the largest viral genome database, IMG/V4, only 501 viruses overlapped with CGVD genomes, emphasizing the high host specificity of chicken enteric phages.

Interestingly, we identified 19.74% (723 out of 3,663) of the CMKMC catalog’s viruses within CGVD, but these accounted for only 1.95% of the total CGVD vOTUs (Fig. 1B). To further expand the CGVD, we incorporated 2,360 medium- or high-quality unique viruses from the CMKMC dataset. This integration brought the CGVD to its current size of 39,380 non-redundant viral genomes, making it the most extensive collection of chicken GI virus genomes available. The integrity of all viral genomes in the CGVD catalog was rigorously evaluated using CheckV. Among these, 19.13% were classified as complete genomes, 26.24% as high-integrity, and 54.63% as medium-integrity genomes (Fig. 1C).

Using protein similarity and gene-sharing metrics, the 39,380 vOTUs were grouped into 3,084 approximate genus-level clusters and 404 approximate family-level clusters. Accumulation curve analysis indicated that the CGVD catalog is nearing saturation at the family level but remains incomplete at the genus level (Fig. 1D-E).

### Taxonomic annotation

Using an updated gene-sharing pipeline [48], 54.03% (21,670 out of 39,380) of vOTUs in the CGVD catalog were successfully assigned to viral families (Table S5). The majority of these vOTUs belonged to three bacteriophage families within the *Caudovirales* order: *Siphoviridae* (n=13,691), *Myoviridae* (n=3,321), and *Podoviridae* (n=1,505).

In addition to these dominant families, other viral families were also identified among the CGVD catalog. These included *Microviridae* (n=699), *Salasmaviridae* (n=526), *Podoviridae_crAss-like* (n=423), *Autographiviridae* (n=286), *Rountreeviridae* (n=251), *Quimbyviridae* (n=205) and *Drexlerviridae* (n=143) (Fig. 2A). These families were frequently observed among the remaining unclassified vOTUs. Interestingly, a significant proportion of the unclassified vOTUs exhibited low completeness levels. Specifically, 60.75% (10,759 out of 17,710) of these unclassified vOTUs had completeness levels below 90%. Interestingly, a significant proportion of the unclassified vOTUs exhibited low completeness levels. Specifically, 60.75% (10,759 out of 17,710) of these unclassified vOTUs had completeness levels below 90%. This observation suggests that these low-completeness vOTUs likely represent novel, yet-to-be-characterized viral genomes. These findings underscore the vast and underexplored diversity of chicken gastrointestinal viruses, which likely exceeds current estimations.

**Figure 2.**
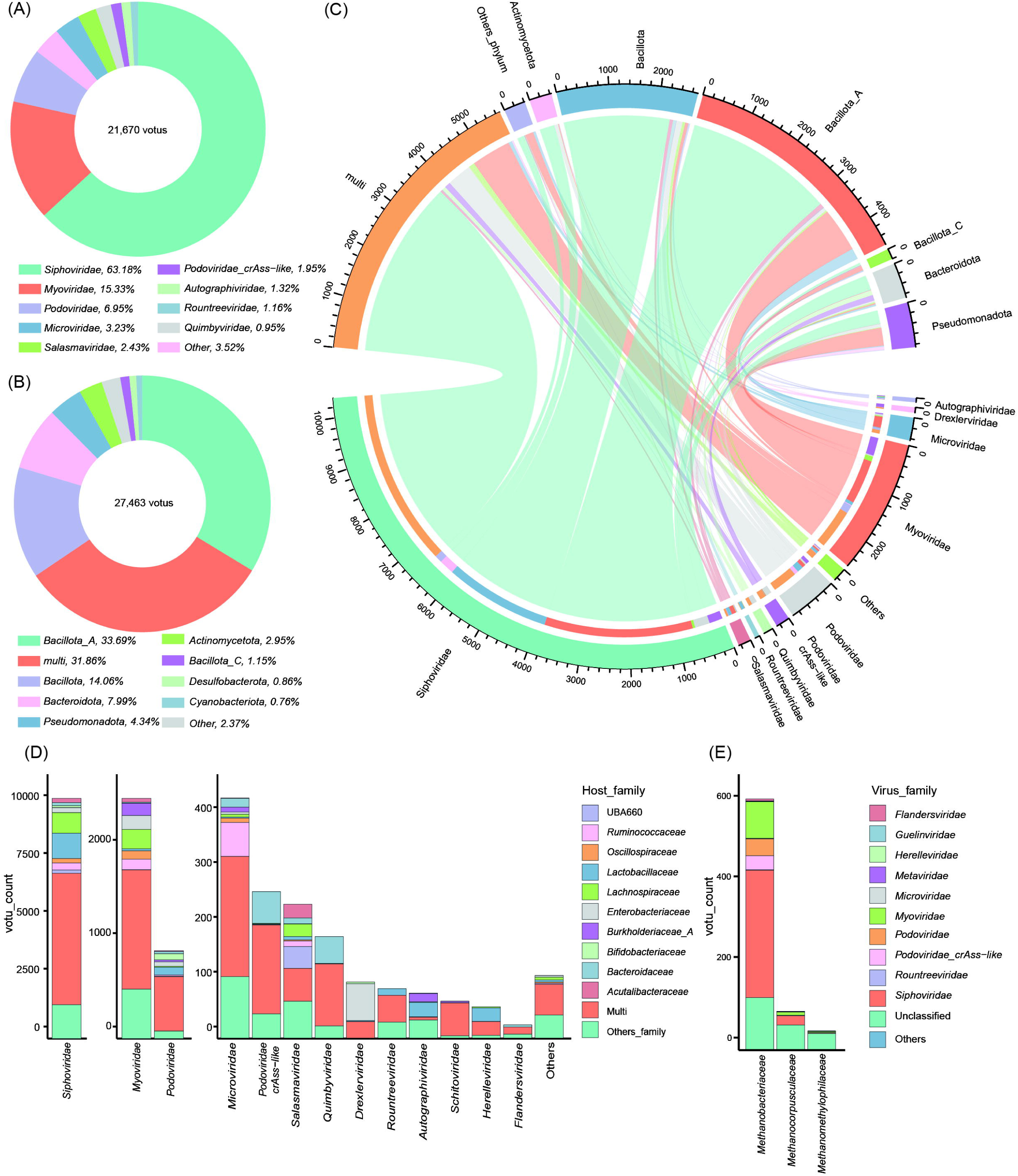
(A) Composition of vOTUs at the family level for known viral classifications. (B) Composition of vOTU hosts at the phylum level for predicted prokaryotic hosts. (C) Quantitative relationship between vOTU family level and host phylum level. (D) Quantitative relationship between vOTU family level and host family level, with the bar chart illustrating the distribution of host family levels for different viral families. (E) Relationship between archaeal family level and viral family level, with the bar chart showing the distribution of viral families across different host families.

### Virus host and lifestyle prediction

We predicted the hosts for the vOTUs by analyzing their homology to genome sequences (>90% nucleotide identity and >30% viral coverage) or CRISPR spacers (bit-score >45) from a large-scale chicken intestinal prokaryotic genome collection comprising 25,827 MAGs (Table S6). This approach enabled host prediction for 69.74% (27,463 out of 39,380) of the vOTUs in the CGVD catalog (Fig. 2B).

At the phylum level, the predicted prokaryotic hosts for chicken intestinal phages were predominantly from *Bacillota_A*, *Bacillota*, *Bacteroidota*, and *Pseudomonadota*, followed by *Actinomycetota, Bacillota_C*, and *Desulfobacterota*. Interestingly, 31.86% (8,749 out of 27,463) of the vOTUs were predicted to infect hosts spanning more than one prokaryotic phylum (Fig. 2C). This proportion is significantly higher than what has been observed in the human gut or oral cavity. Viruses with broad host ranges are generally more capable of spreading and replicating across diverse microbial populations, reflecting strong adaptability. These findings highlight the remarkable diversity and complexity of the viral community in the chicken GI tract.

Within the most abundant viral family in the chicken GI tract, *Siphoviridae*, over half of the vOTUs were predicted to infect at least two bacterial families—a proportion significantly higher than in other viral families. Siphoviridae vOTUs were capable of infecting major bacterial families in the chicken microbiota, including *Lachnospiraceae* and *Lactobacillaceae*. Conversely, *Myoviridae* viruses were predominantly associated with the Proteobacteria phylum, particularly targeting species within *Enterobacteriaceae* and *Burkholderiaceae* (Fig. 2D and Table S7).

Interestingly, 763 vOTUs in the CGVD catalog were predicted to infect archaea, with hosts primarily belonging to *Methanobacteriaceae* and *Methanocorpusculaceae* (Fig. 2E and Table S8). These archaea are involved in anaerobic metabolism and methane production, suggesting that viruses play a role not only in regulating bacterial populations but also in influencing archaeal activity. These findings open up new directions for investigating gas exchange and metabolic processes in the chicken intestine while shedding light on the role of viruses in anaerobic environments.

In addition to host prediction, we performed lifestyle classification of the vOTUs. Among the CGVD catalog, 28.28% (11,137 out of 39,380) of vOTUs were identified as lytic phages with scores >0.5. Of these, 42.17% (4,694 out of 11,137) were predicted to be virulent phages, with scores >0.7 (Table S9). This comprehensive classification provided insights into the host range and ecological roles of viruses within the chicken gut microbiome.

### Phylogenomic analysis of viruses

Using genome-wide protein-level similarity [35], we constructed a proteomic tree for 7,533 complete vOTUs (Fig. 3). Of these, 66.33% (4,997/7,533) were assigned to known viral families, and 66.16% (4,984/7,533) were predicted to infect known prokaryotic hosts. The proteomic tree revealed that viruses clustered based on both family-level taxonomies and potential host affiliations, suggesting that host adaptation plays a critical role in the genomic evolution of chicken GI viruses. Interestingly, *Podoviridae* viruses displayed a unique pattern compared to other families within the *Caudovirales* order. They were predominantly grouped into a single clade in the proteome tree, with most vOTUs infecting bacterial families such as *Lactobacillaceae* and *Bifidobacteriaceae*.

**Figure 3.**
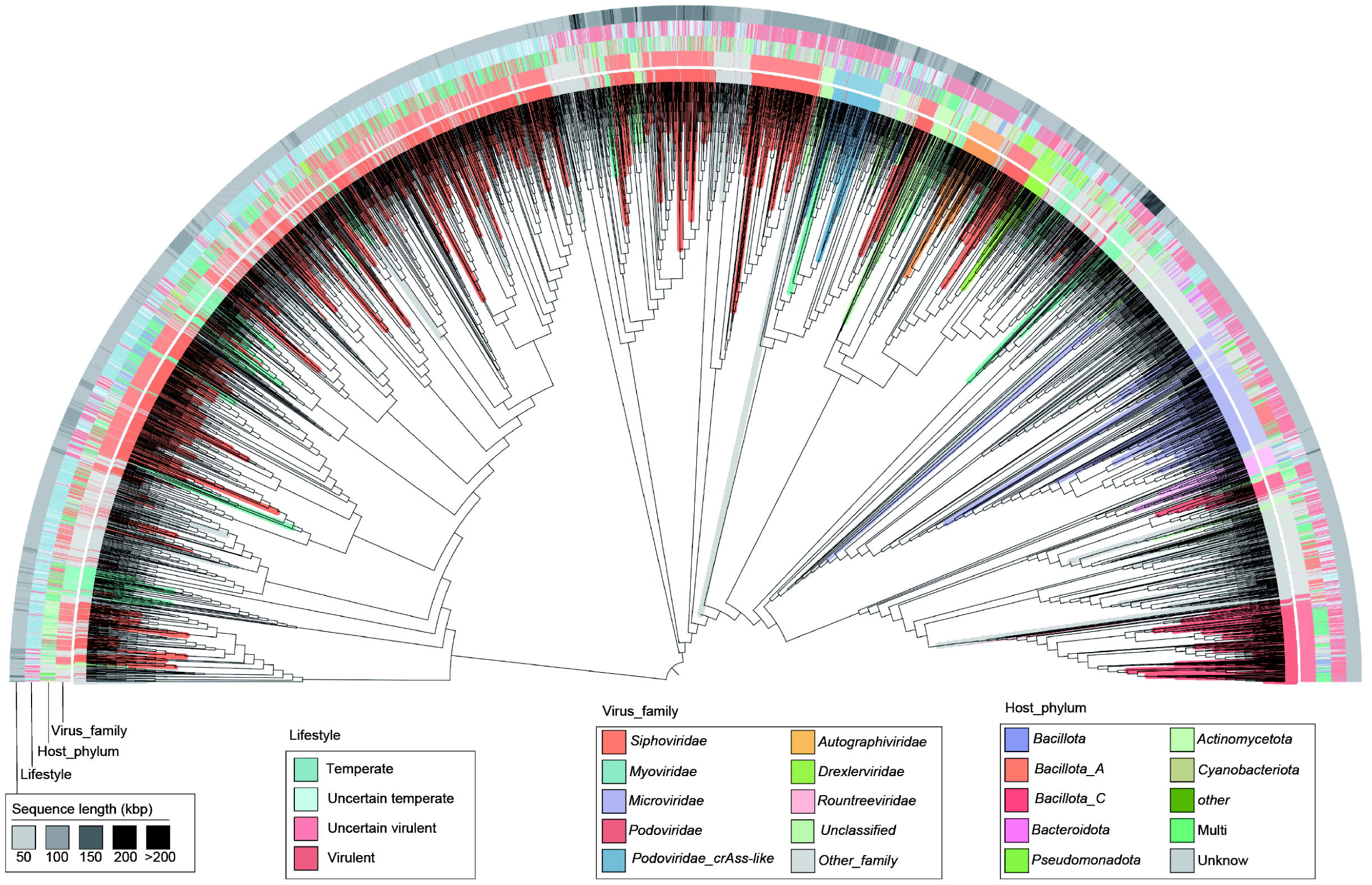
A proteomic tree of 7,233 complete high-quality vOTUs, generated using ViPTreeGen. The tree is annotated with the following metadata: the innermost ring represents viral family-level taxonomic assignments; the second ring shows phylum-level host assignments; the third ring indicates viral lifestyle; and the outermost ring displays the sequence length of each vOTU.

Lifestyle prediction further revealed that most viruses in the *Autographiviridae* family are lytic phages. However, host prediction for this family presented an intriguing finding: while a small subset of *Autographiviridae* vOTUs infected members of *Bacillota*, the majority lacked known host associations. This underscores the complexity of the chicken GI viral community and highlights the potential impact of these viruses on host health, particularly in shaping the microbiome’s structure and function.

We analyzed the genome sizes of *Autographiviridae* phages, which averaged 41.37 kb and ranged from 38.78 kb to 47.67 kb. The moderate genome size of *Autographiviridae* phages suggests a conserved set of essential genes critical for their life cycle and infection processes. This genomic consistency likely explains why these phages cluster together within a single branch of the proteomic evolutionary tree.

### Functional annotation of CGVD

We hypothesized that functional annotation of the CGVD provides significant insights into the ecological roles of viruses, their interactions with host microbiota, and their contributions to gut metabolic processes. To investigate the functional potential of the chicken enterovirome, we identified 2,020,371 protein-coding genes from 39,380 viral sequences of medium or high quality. These genes were functionally annotated using the KEGG, Pfam, and VOG databases. The results showed that 12.31% (248,707/2,020,371) of the viral genes were assigned KEGG ortholog annotations, 22.35% were mapped to the Pfam database, and 34.49% to the VOG database. KEGG annotations revealed that, among these annotated genes, 41.37% were involved in genetic information processing, 36.45% encoded viral AMGs, and 7.27% were associated with cellular processes (Fig. 4A).

**Figure 4.**
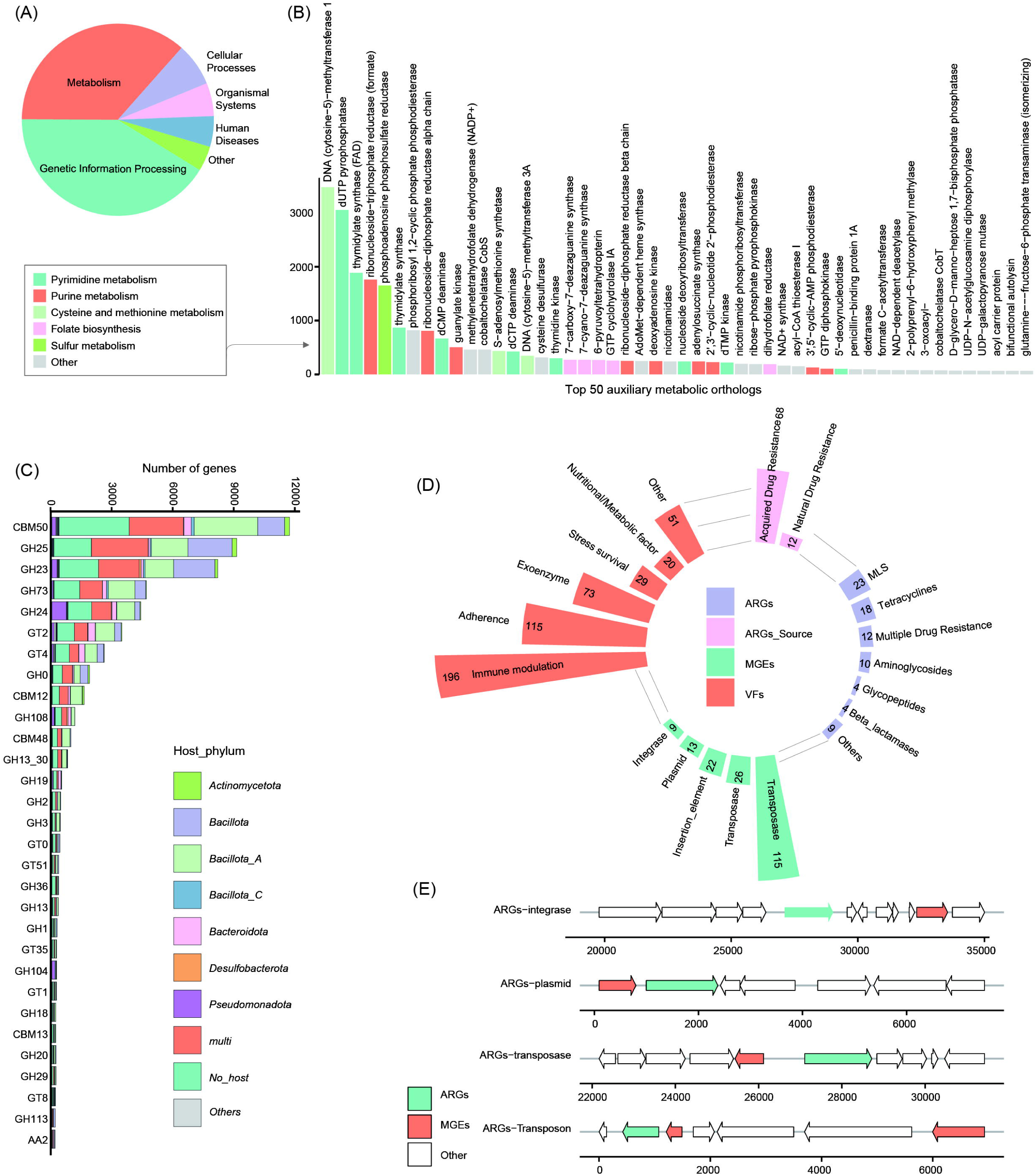
(A) Functional distribution of KEGG-annotated genes in CGVD. (B) Number of genes from the top 50 auxiliary metabolic orthologs in CGVD. An auxiliary metabolic ortholog is defined as a KEGG functional ortholog associated with a KEGG metabolic pathway. (C) Distribution of carbohydrate-active enzymes (CAZymes) encoded by the vOTUs. Enzyme categories include CBM (carbohydrate-binding module), CE (carbohydrate esterase), GH (glycoside hydrolase), GT (glycosyltransferase), and PL (polysaccharide lyase). (D) The number of antibiotic resistance genes (ARGs), virulence factor genes (VFs), and mobile genetic elements (MGEs) in CGVD. (E) Arrow diagrams illustrating specific combinations of ARGs and MGEs within the viral genome. Right arrows denote genes on the forward strand, while left arrows indicate genes on the reverse strand.

To better understand the interaction between the chicken GI virome and the host, we focused our analysis on the AMGs. Mapping these genes to metabolic pathways revealed their involvement in diverse metabolic processes, such as carbohydrate metabolism, amino acid metabolism, and metabolism of cofactors and vitamins. Most AMGs were primarily associated with nucleotide metabolism (46.02%), amino acid metabolism (16.96%), and metabolism of cofactors and vitamins (13.85%)._To investigate the impact of chicken GI viruses on the intestinal microbiome’s biochemical cycles, we further analyzed these AMGs based on KEGG annotations (Table S10). The results showed that 7,771 AMGs were involved in key metabolic pathways, including pyrimidine metabolism, 4,276 in purine metabolism, and 4,258 in cysteine and methionine metabolism (Fig. 4B).

CAZyme annotation revealed that only 2.93% of the protein-coding genes were associated with carbohydrate-active enzymes (CAZymes) (Table S11). These genes primarily encoded enzymes related to peptidoglycan lyase/hydrolase activity (Fig. 4C). Interestingly, 69.39% (27,326/39,380) of vOTUs in CGVD encoded at least one CAZyme. Of these, 72.20% (19,680/27,326) were predicted to infect prokaryotic hosts, suggesting that viral modulation of bacterial cell wall degradation is likely significant during infection.

### Analysis of antibiotic resistance and virulence genes

Using Diamond’s BLASTp tool, we compared viral protein-coding genes with the CARD, MGE, and VFDB databases. Regarding antibiotic resistance genes (ARGs), 80 ARGs were identified in the CARD database, with 12 classified as natural resistance genes and 68 as acquired resistance genes. The most common categories of resistance genes were macrolide-lincosamide-streptogramin (n=23), tetracyclines (n=18), and multidrug resistance (n=12). Only four beta-lactamase resistance genes were detected (Fig. 4D and Table S10).

Considering the key role of MGEs in facilitating the horizontal transfer of ARGs within and between bacterial cells, we assessed the potential for transmission of viral ARGs. Our analysis identified 185 MGEs, classified into three major categories: transposases (n=115), transposons (n=26), and insertion elements (n=22) (Fig. 4D and Table S11). We evaluated the mobility of these ARGs by identifying ARGs located within 5 kilobases of MGEs, as previously established [49], considering MGE-ARG combinations as mobile ARGs. This analysis revealed only 15 vOTUs with MGE-ARG combinations involving four MGE types (Fig. 4E). This suggests that viruses play a limited role in the horizontal transfer of ARGs. Furthermore, we explored the distribution of virulence factor genes (VFGs) in the CGVD vOTUs. Among the CGVD vOTUs, the most common VFGs were involved in immune modulation (n=196) and adherence (n=115) (Fig. 4D and Table S12). This pattern is consistent with findings from previous studies.

### Regional differences in the CGVD

To determine the composition of the enterovirus community, we mapped clean reads from 575 chicken intestine samples to the CGVD database. The average mapping rate was 10.71%, ranging from 0.05% to 34.60% (Fig. 5A), indicating that the database efficiently captured enterovirus reads. As the sample size increased, the dilution curve gradually leveled off, suggesting that the vOTUs in CGVD sufficiently represent the diversity of viruses present in the chicken intestine (Fig. 5B).

**Figure 5.**
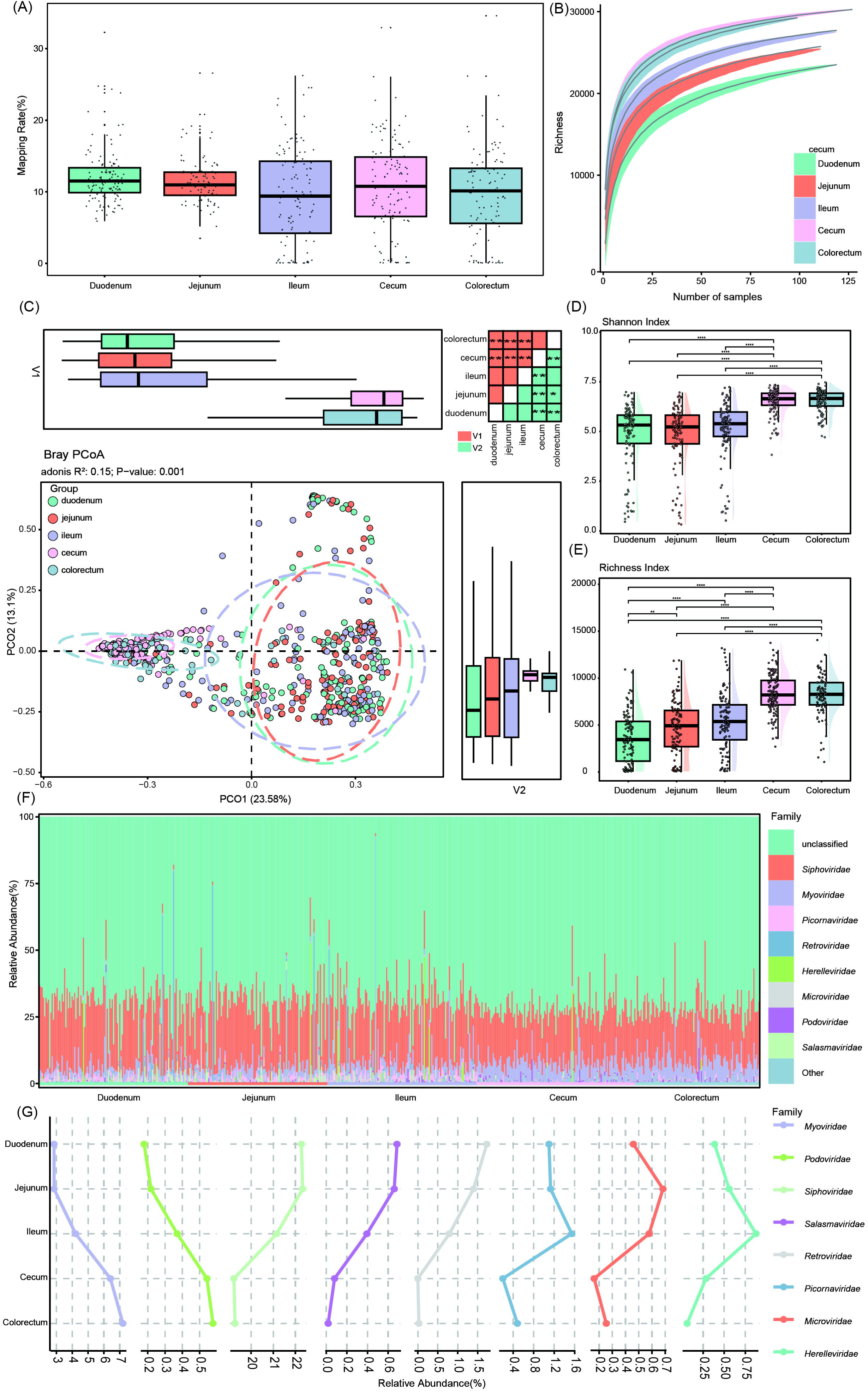
(A) Boxplots illustrating the proportion of metagenomic reads mapped to CGVD in metagenomic data from different regions of the chicken gut. (B) Rarefaction curve analysis showing the relationship between the accumulation of viral genome abundance and the increase in sample size. (C) Principal Coordinates Analysis (PCoA) of Bray-Curtis distances for the enterovirome at the vOTU level. Samples are plotted along the first and second principal coordinates (PCoA1 and PCoA2), with the variance explained by each axis indicated. Ellipses represent 95% confidence intervals around each group. The upper and right boxplots show the fraction of samples along PCoA1 and PCoA2 (boxes represent medians and quartiles; with error bars extending to the most extreme value within 1.5 interquartile ranges). The heatmap in the upper right corner illustrates significant differences between the groups along the first and second axes, as well as between the second and third axes. Statistical significance was determined using the Wilcoxon rank-sum test: \**p* < 0.05, \*\**p* < 0.01, \*\*\**p* < 0.001. (D) Boxplot showing the Shannon diversity index of the enterovirome across all samples. Statistical significance was determined using the Wilcoxon rank-sum test: \**p* < 0.05; \*\**p* < 0.01. (E) Boxplot showing the richness index of the enterovirome across all samples. Statistical significance was determined using Fisher’s test: \**p* < 0.05; \*\**p* < 0.01. (F) Bar graph showing the composition of the enteroviral community (at the family level) in different regions of the chicken intestine. Only the top 8 most abundant viral families are shown. (G) Line graph illustrating the dynamic changes in the relative abundance of the top 8 most abundant viral families across different regions of the chicken intestine.

We conducted PCoA based on the Bray-Curtis distance of vOTU profiles to evaluate compositional differences in the enterovirome across different intestinal regions. The first two principal axes explained 36.68% of the total variation, with clear separation observed between samples from the large intestine (caecum and colorectum) and those from the small intestine (duodenum, jejunum, and ileum). Large intestine samples had significantly higher scores on the first principal coordinate compared to small intestine samples. A similar trend was observed along the second principal coordinate. These differences were statistically significant, as confirmed by PERMANOVA (R² = 0.15, *p* = 0.001) (Fig. 5C).

To assess the richness and evenness of the enterovirome in different intestinal regions, we calculated alpha diversity using richness and Shannon indices. Statistical analyses using Fisher’s method and the Wilcoxon rank-sum test revealed that the duodenum exhibited the lowest diversity and richness, while the colorectum had the highest. Diversity and richness consistently increased from the front (duodenum) to the back (colorectum) of the intestinal tract (Fig. 5D-E). Significant differences in enterovirome composition were also observed between the large and small intestines, highlighting marked regional variations in viral diversity and composition.

At the family level, a substantial portion of viral sequences (mean relative abundance 70.22%) was assigned to vOTUs belonging to unknown viral families, emphasizing the need for further exploration of chicken enteroviruses. Among the identified viral families, *Siphoviridae* (mean relative abundance 20.86%) and *Myoviridae* (mean relative abundance 4.67%) were the most dominant across all intestinal regions. Lower-abundance families included *Picornaviridae* (mean relative abundance 0.89%), *Retroviridae* (0.80%), *Herelleviridae* (0.44%), *Microviridae* (0.43%), *Podoviridae* (0.43%), and *Salasmaviridae* (0.37%) (Fig. 5F).

Regional variations in viral family abundance were also evident. *Siphoviridae*, *Picornaviridae*, *Retroviridae*, *Herelleviridae*, *Microviridae*, and *Salasmaviridae* were predominantly found in the small intestine. In contrast, *Myoviridae* and *Podoviridae* were mainly distributed in the large intestine (Fig. 5G). These findings underscore the marked regional differences in the composition of chicken enteroviruses, reflecting the complexity and diversity of the viral community along the intestinal tract.

## Discussion

The CGVD catalog represents the first comprehensive resource for understanding the viral landscape within chicken gut. It was developed from 3,312 metagenomic datasets spanning 49 studies conducted in 18 countries, making it the largest global dataset for the chicken GI tract to date. This extensive coverage incorporates regional variations in poultry practices, diet, and microbial exposure, ensuring that the findings are ecologically and globally relevant. By integrating high-quality genomes from the CMKMC, the catalog included 39,380 non-redundant viral genomes, balancing comprehensive coverage with accuracy. This has established the CGVD as a robust resource for comparative analyses and functional research.

Taxonomic analysis revealed that the catalog contained 3,084 genus-level and 404 family-level clusters. Although the family-level classification indicated near saturation, the genus-level clustering highlighted the need for further exploration to identify additional diversity. Interestingly, 90% (35,793 of the 39,380) CGVD genomes were unique to the chicken gut, when compared to established databases (e.g., IMG/VR4), demonstrating the distinct nature of the chicken gut virome and its potential for co-evolution with the chicken gut microbiome. The catalog revealed the remarkable taxonomic diversity of the chicken gut virome, encompassing both well-characterized and novel viral families, with the viral communities being predominantly composed of members of the Caudovirales order, including the *Siphoviridae*, *Myoviridae*, and *Podoviridae* families. These families also dominate human gut virome catalogs [50]. These findings align with observations from gut viromes in honeybees, birds, lizards, and mammals [51-53].

More than half of the vOTUs were assigned to established families, including *Siphoviridae*, *Myoviridae*, and *Podoviridae*, which play key ecological roles by regulating microbial dynamics, facilitating transduction of genetic material in bacterial communities, and encoding auxiliary metabolic genes that may influence nutrient absorption, disease resistance, the phenotype of the bacterial hosts, the gut health and functionality [54-56]. Less-represented families, such as *Microviridae* and *Quimbyviridae*, along with newly proposed families such as *Drexlerviridae*, added complexity to the virome’s ecological landscape and underscored the potential for novel discoveries. The identification of *Podoviridae_crAss*-like viruses associated with *Bacteroides* indicated evolutionary links between chicken and human gut viromes, raising considerations about cross-host microbiome interactions and zoonotic transmission [57, 58]. The catalog also revealed a substantial proportion of unclassified vOTUs, pointing to gaps in existing viral taxonomies and emphasizing the need for better sequencing technologies and more robust classification frameworks. Many of these unclassified vOTUs showed low genome completeness, indicating that they likely represent novel viral lineages requiring further characterization.

The results provided novel insights into host-virus interactions, revealing the intricate roles of viruses in shaping microbial communities and regulating gut ecosystem dynamics. Host predictions were successfully made for 69.74% of vOTUs, demonstrating the robustness of the CRISPR spacer homology approach and the quality of the associated prokaryotic genome collection. About 31.86% of vOTUs displayed broad host-range adaptability, infecting multiple prokaryotic phyla and positioning these viruses as key regulators for stabilizing microbial communities and promoting gut resilience. Specific host associations were also observed, with *Siphoviridae* phages promoting beneficial bacterial families, *Lachnospiraceae* and *Lactobacillaceae*, supporting nutrient metabolism, while *Myoviridae* phages targeted pathogens such as *Proteobacteria*, indicating their dual roles in maintaining gut homeostasis and controlling harmful bacteria.

The results reveal the impact of the chicken gut virome on fermentation, gas production and energy flow. A significant discovery was the identification of 763 vOTUs targeting archaea, primarily *Methanobacteriaceae* and *Methanocorpusculaceae*, which play key roles in methane production. This highlighs the potential for phages to influence archaeal populations and suggests innovative strategies for mitigating greenhouse gas emissions, leveraging virome research to enhance environmental sustainability in poultry farming. Such an approach highlights the intersection of virome research with global sustainability efforts, positioning it as a valuable tool in addressing climate change. In addition to these environmental implications, 28.28% of vOTUs were classified as lytic phages, known for their roles in microbial turnover, nutrient recycling, and pathogen control [59, 60]. The adaptability and diversity of chicken gut phages, along with their regulation of bacterial and archaeal taxa, highlight their importance in microbial management and sustainable farming practices.

The phylogenomic analysis of the CGVD catalog highlighted host specificity and adaptation as key drivers of viral evolution, evident in distinct phylogenomic patterns observed in families such as *Podoviridae* and *Autographiviridae*. 66.33% of the vOTUs were assigned to known viral families, and 66.16% were linked to predicted hosts, underscoring the significant influence of host adaptation on viral diversification. For instance, *Podoviridae* phages displayed a strong preference for bacterial families *Lactobacillaceae* and *Bifidobacteriaceae*, suggesting a co-evolutionary relationship that contributes to microbiome stability and the maintenance of beneficial bacterial populations.

Lifestyle prediction analysis of *Autographiviridae* phages revealed their role as lytic phages, driving bacterial population control. These phages exhibited evolutionary conservation in genome size (averaging 41.37 kb), reflecting specialization for efficient replication within specific ecological niches. Genome size directly impacts the diversity and functionality of encoded genes [61], highlighting the evolutionary trade-offs in phage adaptation. The presence of unclassified vOTUs points to gaps in understanding host-virus interactions, suggesting the existence of novel hosts or host-switching mechanisms that could expand our knowledge of virome complexity.

Functional annotation analyses of AMGs and CAZymes highlighted their significant roles in regulating microbial metabolism, nutrient cycling, microbial turnover, virus-environment interactions, aligning with prior research on viral contributions to the ecology and evolution of bacterial communities [62]. From 39,380 viral sequences, 2,020,371 protein-coding genes were identified, with notable proportions linked to KEGG orthologs (12.31%), Pfam (22.35%), and VOG (34.49%). However, most genes remain unannotated, underscoring the novelty of these viral genomes and the need for expanded genomic resources to characterize viral diversity and discover the functions of new genes. The analysis revealed that 41.37% of KEGG orthologs were associated with genetic information processing, highlighting viruses’ dependence on host mechanisms for DNA/RNA synthesis, repair, and translation—processes critical for replication. This aligns with the well-established concept that viral auxiliary genes support viral replication by promoting the degradation of host DNA and RNA and redirecting host metabolism toward nucleotide biosynthesis [63].

AMGs, comprising 36.45% of the identified KEGG orthologs, play critical roles in viral adaptation by co-opting host metabolic pathways to enhance replication efficiency. These genes were linked to host nucleotide metabolism (46.02%), amino acid metabolism (16.96%), and cofactors and vitamins metabolism (13.85%), highlighting their influence on microbial dynamics and ecosystem functions. Specifically, AMGs involved in purine and pyrimidine metabolism support nucleic acid synthesis, directly facilitating viral genome replication [63]. Those associated with cysteine and methionine metabolism may enhance antioxidant responses, mitigating oxidative stress and promoting protein synthesis, which benefits both viral and bacterial survival in the gut [64]. These findings underscore the role of viruses in modulating host biochemical pathways to enhance replication, adapt to host environments, and potentially influence host immune responses [65].

The functional profiles of CGVD viruses shared similarities with human gut viromes, particularly in AMGs related to nucleotide metabolism. Considerable phage-associated functional genes contribute to the adaptive evolution of their hosts, including genes involved in nutrient metabolism, energy metabolism, DNA repair and virulence [66-69]. In our study, the higher prevalence of CAZymes suggests unique adaptations to chicken gut physiology and diet, emphasizing the importance of species-specific virome research to understand host-specific viral strategies and their ecological impacts. The identification of CAZymes, present in 69.39% of vOTUs, involved in peptidoglycan degradation, underscores the role of viruses in bacterial cell wall lysis [70], highlighting the impact of viruses on microbial turnover, microbial competition, nutrient cycling, and gut stability. These findings highlight the role of viruses as active modulators of host metabolism and gut microbiota, with implications for optimizing poultry health, productivity, and pathogen resistance.

The CGVD catalog provided valuable insights the contribution of the of chicken gut viruses to ARG and VFG dynamics, as well as regional variations in virome composition and diversity across the GI tract. Bacteriophages can contribute to the dissemination of ARGs through horizontal gene transfer [71]. Phage-associated ARGs have been discovered from various sources that are closely related to human activities, such as animals, including livestock and poultry [72]. A high abundance of macrolide-lincosamide-streptogramin resistance genes has been detected in the cecum microbiome of chicken [73]. In our study, we identified 80 antimicrobial resistance genes (ARGs), with the majority linked to resistance mechanisms such as macrolide-lincosamide-streptogramin resistance, tetracycline resistance, multidrug resistance, and beta-lactamase production, suggesting that viruses play a secondary role in the transfer of ARGs compared to bacterial plasmids and integrative elements. The relatively low abundance of ARGs in viral genomes, when compared to bacterial hosts suggests a limited role for viruses in the propagation of ARGs. The scarcity of mobile genetic elements within viral genomes further reinforces this result. However, the detection of both natural and acquired ARGs within the virome suggests that viral genomes may act as reservoirs for ARGs, thereby indirectly influencing the spread of antibiotic resistance in microbial communities [74, 75]. This highlights the potential for viruses to contribute, through transduction or other mechanisms, to the horizontal transfer of ARGs across different hosts and environments [76-79].In addition, the catalog included 311 VFGs, predominantly associated with immune modulation and bacterial adhesion. This underscores the role of viruses in modulating bacterial pathogenicity and their potential impact on host-microbiome interactions [80, 81].

This study underscores the regional specificity of the chicken enterovirome, revealing significant differences in the composition and abundance of enteric viruses across intestinal regions. Similar findings have been reported in the domestic pig and rhesus macaque [82] and humans [83]. Alpha diversity analysis demonstrated a progressive increase in viral diversity from the small intestine to the large intestine, likely driven by the accumulation of diverse microbial populations in the distal regions [84]. This gradient may be influenced by variations in immune responses, pH levels, nutrient availability, and other factors that create distinct ecological niches favoring specific viral families [85-88]. The increasing complexity of the virome along the GI tract reflects dynamic interactions between viruses, bacteria, and other microbial inhabitants, which collectively shape the community structure of viruses. Principal coordinate analysis further revealed distinct viral community profiles between the small and large intestines. Viral families such as *Siphoviridae* and *Picornaviridae* were more prevalent in the small intestine, while *Myoviridae* and *Podoviridae* dominated the large intestine. These differences likely correspond to the unique physiological and microbial environments of each intestinal region. For instance, the small intestine is primarily involved in nutrient absorption and harbors a relatively lower microbial density [89], which may favor the persistence of certain viral taxa. In contrast, the large intestine serves as a hub for microbial fermentation and metabolism [90], supporting a more diverse and metabolically active microbial ecosystem that may influence the composition of the virome.

## Conclusions

The CGVD catalog provides a comprehensive and valuable resource for understanding the chicken gut virome, offering insights into its ecological, evolutionary, and functional roles. The catalog revealed the crucial role of phages in regulating microbial dynamics, supporting gut health, and controlling pathogens. The CGVD catalog not only advances our understanding of host-virus interactions but also offers practical applications for sustainable poultry health management, including the development of precision phage therapies and reduced reliance on antibiotics. In addition, this study provides significant insights into the regional dynamics of the chicken gastrointestinal virome, offering a comprehensive resource for understanding viral diversity, community structures, and their functional roles in poultry health. These findings lay a robust foundation for future investigations into the interplay between viruses and their microbial and host environments, potentially informing strategies to enhance poultry health through targeted virome modulation.

## Declarations

### Ethics approval and consent to participate

Not applicable

### Consent for publication

All authors have seen and approved the final, submitted version of this manuscript.

### Availability of data and material

The project numbers of all metagenomic samples involved in this study are provided in Additional file 1. The CGVD dataset constructed in this study has been deposited in the Zenodo repository under the accession number 14684892 (doi.org/10.5281/zenodo.14684892).

### Authors’ contributions

J.X.Z: Writing– original draft, Visualization, Methodology, Investigation, Formal analysis, Data curation. H.B.N: Writing – Review & Editing, Data curation, Funding Acquisition. F.L.N: Writing – Review & Editing, Methodology, Visualization, Formal analysis. Q.Y and S.L: Writing – Review & Editing, Methodology, Visualization, Software. Y.C, J.X.M and H.L.Y: Writing – Review & Editing, Formal analysis. K.M.S, R.L and X.W.H: Writing – Review & Editing, Data curation. H.M.E and H.M: Conceptualization, Writing – Review & Editing, Supervision. X.X.Z: Conceptualization, Writing – Review & Editing, Project Administration, Resources, Supervision.

## Acknowledgements

Not applicable

## Supplementary Materials

**Additional file 1: Chicken metagenomic data and sample information. Tables S1**: Details of the chicken metagenomic data. **Tables S2**: Sample information from different regions of the chicken intestine. https://doi.org/10.5281/zenodo.15224168

**Additional file 2: Quality assessment of viral genomes. Tables S3**: Quality assessment of all vOTUs with medium quality or higher before clustering. **Tables S4**: Quality assessment of viruses in the CGVD. https://doi.org/10.5281/zenodo.15224168

**Additional file 3**: **CGVD taxonomic and host predictions**. **Tables S5**: Family-level species annotation information in the CGVD. **Tables S6**: Phylum-level and family-level host prediction information in the CGVD. **Tables S7**: Lifestyle prediction results in the CGVD. https://doi.org/10.5281/zenodo.15224168

**Additional file 4: Functional and genetic analysis of viruses**. **Tables S8**: Details of auxiliary metabolic genes. **Tables S9**: CAZyme database annotation results; **Tables S10**: Drug resistance gene annotation results. **Tables S11**: Mobile genetic element annotation results. **Tables S12**: Virulence factor annotation results. https://doi.org/10.5281/zenodo.15224168

